# Thermal preconditioning in a reef-building coral alleviates oxidative damage through a BI-1 mediated antioxidant response

**DOI:** 10.1101/2021.03.15.435543

**Authors:** Eva Majerová, Crawford Drury

## Abstract

Global coral reef decline is largely driven by the breakdown of the coral-algal symbiosis during temperature stress. Corals can acclimatize to higher temperatures, but the cellular processes underlying this ability are poorly understood. We show that preconditioning-based improvements in thermal tolerance in *Pocillopora acuta* are accompanied by increases in host glutathione reductase (GR) activity and expression, which prevents DNA damage. A strong correlation between *GR* and *BI-1* expression in heat-stressed preconditioned corals and the presence of an antioxidant response element (ARE) in the *GR* promoter suggest BI-1 could regulate *GR* expression through Nrf2/ARE pathway. To fortify this link, we developed an siRNA-mediated gene knockdown protocol and targeted the coral BI-1 gene. BI-1 knock-down specifically decreased GR expression and activity and increased oxidative DNA damage in heat-stressed preconditioned corals, showing that BI-1-mediated enhanced antioxidant response during acute heat stress is a key mechanism that prevents oxidative DNA damage after preconditioning.

**Teaser:** Preconditioning improves redox homeostasis and prevents oxidative stress in a thermally stressed reef-building coral

## Introduction

Healthy coral reefs support nearly one-third of marine species and provide shelter, nursery habitat and coastal protection across the tropical oceans ^1–3^. The energetic and structural foundation of this ecosystem is the coral-algal symbiosis, which is disrupted by thermal stress and is critically threatened by climate change. Climate induced mass bleaching has impacted most of the world’s reefs and is predicted to increase in frequency and intensity, threatening the long-term persistence of these ecosystems ^4–6^. A deeper understanding of the cellular and molecular mechanisms underlying coral-algal symbiosis maintenance is critical for modern conservation and management ^7, 8^.

Corals are energetically dependent on their intracellular algal symbionts (family *Symbiodiniaceae*)^9^, which provide photosynthetically synthesized sugars and receive shelter and a supply of inorganic molecules from the host. During thermal stress, symbionts release increased reactive oxygen species (ROS), which are believed to trigger molecular cascades resulting in coral bleaching and can lead to the eventual death of the coral host organism ^10–18^; however, definitive proofs for this hypothesis are still missing ^19^.

Under normal conditions, ROS are scavenged by antioxidant systems in the host and symbionts to reduce damage to cell membranes, lipids and nucleic acids ^20, 21^. Under increased temperatures, ROS concentrations become elevated and the coral holobiont activates first-line enzymatic antioxidants such as catalase or superoxide dismutase and maintain a reducing intracellular environment via the glutathione redox cycle ^11, 13, 15, 17, 22–25^. These systems are well-studied in *Symbiodiniaceae*, where genera with different thermal resilience vary in antioxidant gene expression ^13, 17, 24^ and thermally resilient algae generally produce more antioxidants, which are able to better maintain cellular homeostasis. Conversely, the role of host-derived antioxidants in bleaching and thermal resilience remains disputed. While pioneering studies show antioxidant activation in adult coral host tissue and larvae under heat stress ^17, 26–30^, more recent experiments have failed to find this pattern ^11, 23^. However, the coral host control of the level of oxygen radicals and ROS is likely a major component in the dynamics of coral-algal symbiosis maintenance ^19, 31^ and activation of DNA damage response pathways is a hallmark of thermal stress in corals^32^

Intra-generational plasticity is critical for mitigating stress in corals, which have a limited ability to avoid stressful environmental conditions, highlighting the importance of acclimatization for the long-term persistence of coral reefs by ‘buying time’ for adaptive change ^33^. While many corals show improved thermal tolerance after pre-exposure to sublethal temperatures ^34–41^, the molecular triggers and consequences of this process remain poorly understood.

In this work we show that thermal preconditioning in the cosmopolitan reef-building coral *Pocillopora acuta* prevents heat-induced oxidative DNA damage via increased activity of glutathione reductase. This glutathione reductase activity is highly correlated with coral BI-1 (Bax-inhibitor 1) gene expression in heat-stressed corals. To fortify this link, we developed, validated and conducted siRNA-mediated gene knockdown experiments in living adult corals. First, we validated the method downregulating the coral family of green fluorescent proteins (GFP) proving that the assay leads to both knock-down of GFP gene expression and decrease in GFP autofluorescence signal in host tissue. We then show that BI-1 downregulation leads to a decreased glutathione reductase expression and activity and results in an accumulation of oxidative DNA damage in coral tissue upon acute heat stress. We also used evidence from vertebrate models to document an antioxidant response element (ARE) in a putative promoter region of the coral glutathione reductase gene, suggesting that BI-1 likely impacts the coral antioxidant system through the Nrf2/ARE signaling pathway.

## Results

### Thermal preconditioning improves bleaching susceptibility in P. acuta

This study uses samples from the coral colonies described in the preconditioning experiment of Majerova et al. ^41^, which were pre-exposed to a 3 day sublethal stress at 29°C (PC) or 26°C (NPC/control) two weeks before exposure to acute heat stress (32°C; Fig. 1A). During this acute stress, preconditioned (PC) corals showed substantially increased symbiosis stability when compared to non-preconditioned (NPC) corals (Fig. 1B). After 3 days of heat stress, NPC corals were visibly bleached while PC corals resembled control, non-treated corals. This difference was more pronounced after 5 days of heat stress. PC corals lost symbionts, but significantly more slowly than in NPC corals (p< 0.001) ^41^.

**Figure 1.**
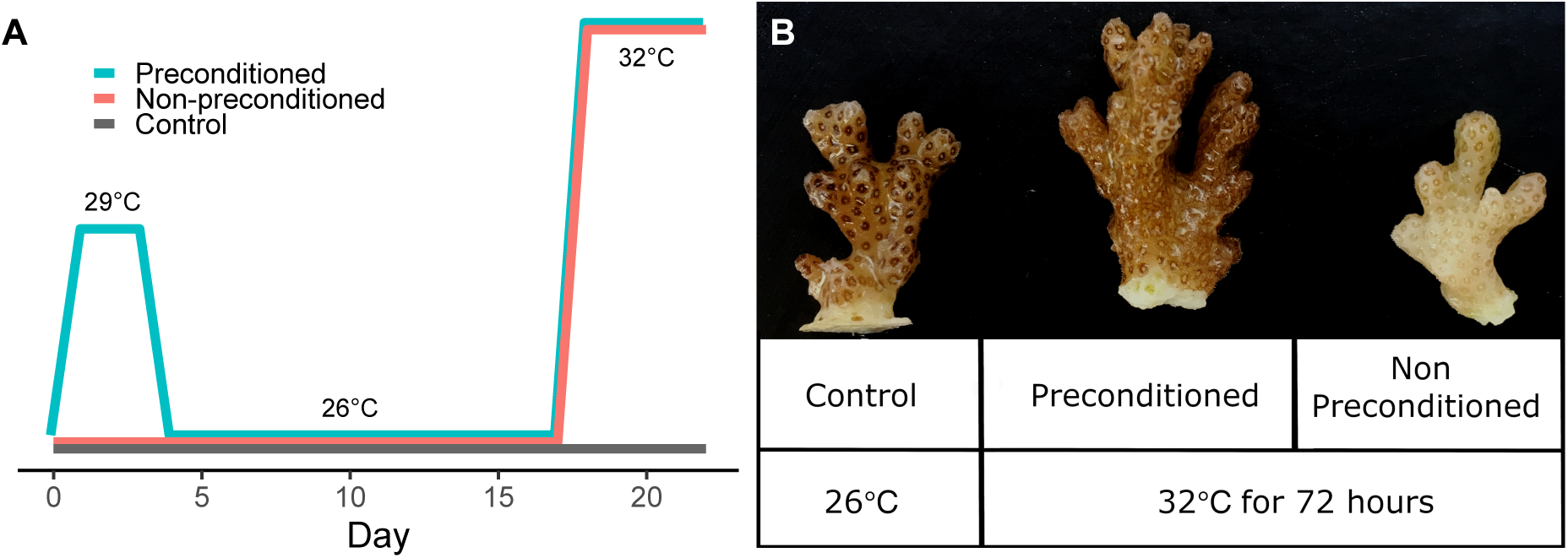
Experimental design. A) Preconditioning profile. Preconditioned corals (PC) were exposed to sublethal 29°C for 72 hours and returned to 26°C for additional two weeks before undergoing acute thermal stress (32°C). Non-preconditioned (NPC) corals were exposed to acute thermal stress with no previous preconditioning. B) Different response to acute heat stress (32°C for 72 hours) in preconditioned (PC) and non-preconditioned (NPC) corals (right). While PC corals resembled control corals, NPC corals were visibly bleached.

### Preconditioning selectively increases the activity of host-derived glutathione reductase

The activity of glutathione reductase in *P. acuta* is influenced by acute heat stress (LMM, activity∼treatment*time + (1|coral); p(time) = 0.001) and was constitutively higher in PC corals than NPC corals (p_(treatment)_ < 0.001, Fig. 2A). Interestingly, the activity of peroxide-scavenging antioxidants was dynamic (p_(time)_ = 0.009) but did not differ between PC and NPC corals. Increased activities of two different peroxidases - catalase and glutathione peroxidase – had been previously detected in stressed corals ^17, 22, 26, 28^, but the catalase activity kit used here is not specific and may detect activity of any enzyme with peroxidase activity (Bioassay Systems, personal communication). Thus, we separately tested the activity of glutathione peroxidase with a specific kit (Bioassay Systems). We were not able to detect any glutathione peroxidase activity in our samples so we conclude that the peroxidase activity is primarily catalase, although we cannot conclusively exclude technical issues with this assay.

**Figure 2.**
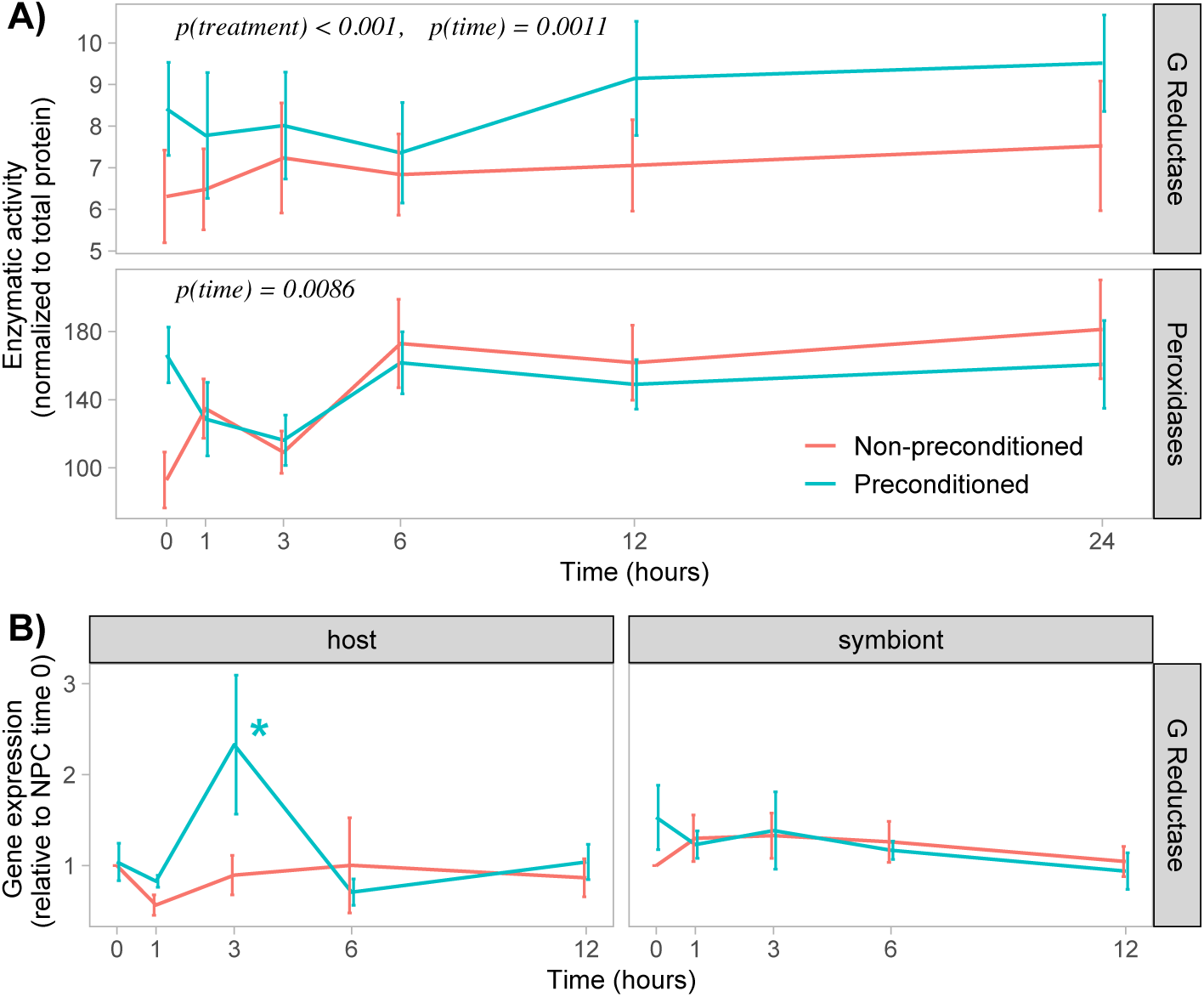
Enzymatic activity and gene expression during thermal stress. A) Thermally *PC* corals *have higher* activity of glutathione *reductase than NPC,* but do not differ in the activity of peroxide-scavenging enzymes. Activity was measured in host cell extracts. *B)* Gene expression pattern of glutathione reductase in host and symbiont cells. Gene expression of host-derived glutathione reductase significantly increased at 3 hours of acute heat stress in *PC corals, but not NPC* corals. Graphs depict means with standard errors, n=6.

To examine if the observed increase in glutathione reductase activity was connected to the stimulation of gene expression, and if such response is common for both partners or is host-specific, we analyzed the expression of coral host and symbiont gene in PC and NPC corals upon heat stress (Fig. 2B). Host glutathione reductase expression changed over time and between treatments (LMM, expression∼treatment*time + (1|coral), p_(time)_ < 0.001, p_(treatment)_ = 0.0098). We observed an increase in expression in PC corals shortly after the beginning of the heat stress (p_(1h)_ = 0.0736) that peaked at 3 hours, when the expression was ∼ 2-fold higher compared to NPC corals (p_(3h)_ = 0.0015).

Symbiont glutathione reductase expression was not significantly different between treatments (p = 0.8099) and did not change over time (p = 0.1485). This suggests that glutathione reductase dynamics are driven by the coral host; however, there was no significant correlation between protein activity and gene expression for either host or symbiont cells (Fig S1), which is a common discrepancy described across the animal kingdom (reviewed in ^42^).

### Increased antioxidant activity protects DNA from oxidative damage

Glutathione reductase helps stabilize the reducing environment of the cell, enhancing its ROS scavenging ability and preventing cellular stress such as oxidative DNA damage ^20, 43^. To clarify whether the increase in glutathione reductase activity improves ROS protection in heat-stressed corals, we analyzed the level of oxidized guanine species (8-OHdG), markers of oxidative DNA damage ^20^. There was a clear difference between PC and NPC corals in time (Two-Way ANOVA, 8-OHdG∼conditioning*time with Tukey post-hoc testing, p_(time:conditioning)_ = 0.0098, Fig. 3). Unlike in PC corals, we observed an accumulation of 8-OHdG in NPC corals after 24 hours of the acute heat stress (p_(NPC)_ = 0.0991, p_(PC)_ = 0.4082, p_(NPC-PC, 24h)_ = 0.0310, p_(NPC – PC, 0h)_ = 0.7449).

**Figure 3.**
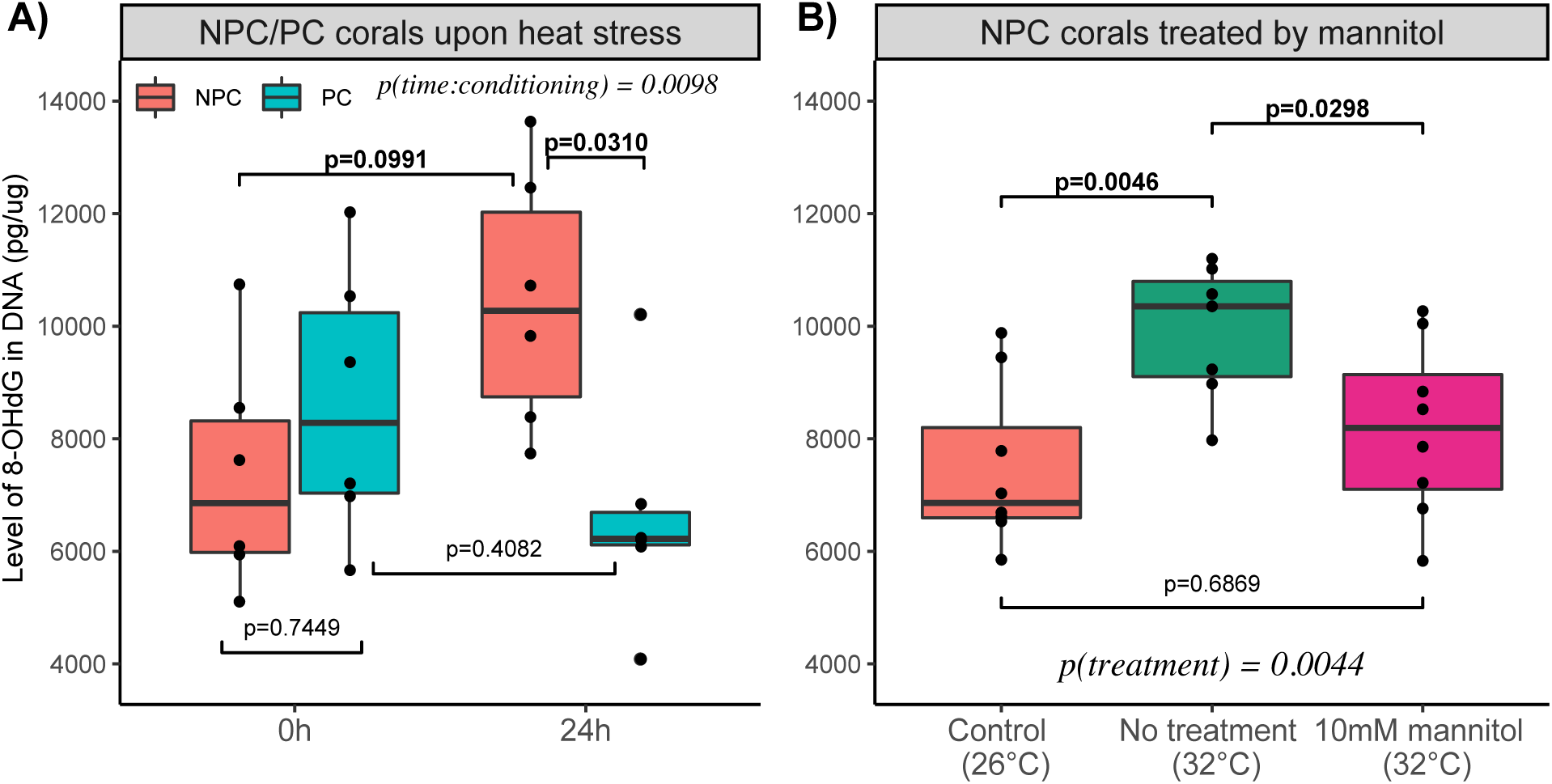
Oxidative DNA damage during heat stress. A) Non-preconditioned corals accumulate DNA damage (the oxidized guanine species, 8-OHdG) but preconditioned corals do not. After 24 hours of the stress, there is a significant difference in the level of 8-OHdG between the treatments. n = 6 B) The accumulation of 8-OHdG in non-preconditioned corals during acute heat stress is prevented by the addition of 10 mM mannitol, a non-enzymatic antioxidant. n = 8. The boxplots show median of the data, first and third quartile and respective datapoints.

To fortify the connection between antioxidants and oxidative DNA damage in corals, we treated NPC corals with 10mM mannitol during acute heat stress. Mannitol is considered a non-enzymatic antioxidant ^44^ that protects plants and algae from ROS and prevents general DNA damage in coral tissue aggregates ^21^. Again, NPC corals accumulated marks of oxidative DNA damage during heat stress (One-way ANOVA, p_(treatment)_ = 0.0044), but the addition of 10mM mannitol eliminated this increase (One-Way ANOVA, p_(no treatment:mannitol)_ = 0.0298, p_(control:mannitol)_ = 0.6869), demonstrating the direct connection between antioxidant defense systems and the DNA damage used here.

pa-BI-1 expression correlates with the expression of glutathione reductase in preconditioned corals.

BI-1 (BAX inhibitor 1) is an anti-apoptotic protein that – among others – promotes cell survival by increasing the production of antioxidants through the activation of Nrf2 transcription factor in human cells ^45, 46^. We previously showed that PC corals increase the expression of pa-*BI- 1* during acute heat stress compared to NPC corals ^41^. Since these observations were made on the same set of samples as our measurements of the expression of glutathione reductase (pa- *GR*), we examined the correlation between the gene expressions of pa-*BI-1* and pa-*GR*. We observed a strong positive correlation in PC (R = 0.948, p_(lm)_ = 0.004) but not NPC (R = 0.126, p_(lm)_ = 0.4679) corals (Fig. S2). The strongest correlation occurred during the first 3 hours of the heat stress, where we observed a significant overexpression of both genes in PC but not NPC corals (^41^ and Fig. 2), suggesting coral BI-1 can regulate the expression of antioxidant genes upon stress conditions.

### siRNA-mediated knockdown assay downregulates coral GFP family

We developed and validated a siRNA-mediated gene knockdown method in living adult corals by targeting proteins of the GFP family, using a common approach for developing similar technologies in non-model organisms^47–49^.

We identified 12 predicted *Pocillopora* GFP-like fluorescent chromoproteins in the NCBI mRNA database and used BLOCK-iT™ RNAi Designer (ThermoFisher Scientific) to design siRNA molecules with the best predicted success rate to inhibit expression of individual predicted GFP-like genes. We chose an siRNA duplex that had 100% sequence similarity with 4 predicted GFP-like genes, one mismatched nucleotide with 7 predicted GFP-like genes and was not complementary to 1 of the genes. In a similar way, we designed RT-qPCR primers to amplify the siRNA target region of the GFP-like genes. Both primers have either full complementarity to the predicted genes or bear one mismatched nucleotide in the central sequence of the primer. We optimized the timing of siRNA administration and coral sampling and used a control siRNA with no known target in *Pocillopora* transcriptome (siNTC) to exclude the effect of the siRNA treatment itself on the studied genes and pathways. We tested several delivery methods, including PEI (Sigma Aldrich), Lipofectamine2000 (Thermofisher), CaCl_2_ phosphate transfection, and Fuse-It (Ibidi), and found the INTERFERin (Polyplus) to be the only one to produce reliable knockdown of the target genes. A total of 16 coral fragments from 8 different parent colonies were used for the siRNA developmental experiment. We analyzed GFP expression and fluorescence signals derived from host GFP and symbiont chlorophyl a (ChlA) for each fragment independently. We found 9 corals (representing 6 out of 8 parent colonies) in which the expression of GFP decreased by at least 20 % compared to the siNTC (Fig. 4A). Expression downregulation ranged from 14% to 58% and averaged 32% ± 16% (mean ± SD). Independent verification of GFP expression via confocal microscopy confirmed the GFP to ChlA ratio decreased to an average of 73% ± 30% (mean ± SD; (p_(T-test)_ = 0.0094) and ranged from 39% to 80% (Fig. 4B). Two corals showed an exception to this pattern where the GFP to ChlA ratio slightly increased despite the expression downregulation.

**Figure 4.**
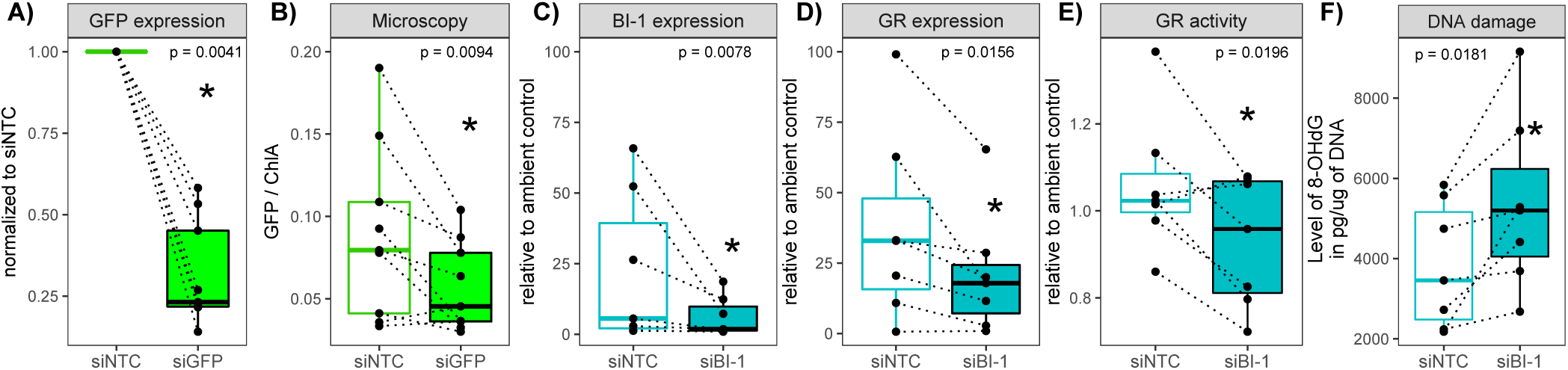
BI-1 overexpression in heat-stressed preconditioned corals improves DNA damage protection via regulation of glutathione reductase activity. A) Expression of GFP in corals treated with siGPF, normalized to siNTC, B) Confocal microscopy analysis of GFP-derived signal normalized to symbiont-derived signal in the same corals, n = 9 C,D) Expression of BI-1 and glutathione reductase genes, respectively, in heat-stressed PC corals is efficiently reduced after siRNA-mediated gene knockdown (siBI-1). siNTC represents PC corals treated with a negative control siRNA. Data are normalized to untreated corals at ambient temperature, n=7 E) The activity of glutathione reductase decreases upon siBI-1 knockdown in PC corals. Data are normalized to untreated corals at ambient temperature, n=7. F) The level of oxidized guanine in coral DNA during heat stress increases after siBI-1 knockdown in PC corals (n = 7). Dotted lines connect paired samples from the same colony. The boxplots show median of the data, first and third quartile and respective datapoints.

### pa-BI-1 controls the expression of glutathione reductase in preconditioned corals

To confirm the presumed link between pa-BI-1 and pa-GR, we inhibited the expression of pa-*BI- 1* in heat-stressed PC corals (Fig. 4C). We optimized the start of the heat stress to 48 hours post-transfection to achieve the strongest knock-down during the first hours of the acute heat stress when *pa-BI-1* was most strongly overexpressed in PC corals ^41^. We successfully inhibited pa-*BI-1* expression in 7 out of 17 corals (similarly to siGPF, we set a threshold of 80% gene expression as a successful knock-down) ranging from 14% to 80% expression compared to siNTC (47% ± 24, mean ± SD; Fig. 4C). For all further analyses, we chose only the 7 corals with a successful knock- down and disregarded the corals with no pa-*BI-1* knock-down.

As expected, we observed an overexpression of both pa-*BI-1* and pa-*GR* in heat-stressed PC corals incubated with siNTC (Fig. 4C,D, siNTC) when compared to control corals (PC corals at ambient temperature). After siBI-1 knockdown (Fig. 4C, siBI-1), the expression of both pa-*BI-1* and pa-*GR* decreased (Fig. 4C,D, Wilcoxon test, p_(BI-1)_ = 0.0078, p_(GR)_ = 0.0156), but was still higher than in ambient control corals (Wilcoxon test, p_(BI-1)_ = 0.0781, p_(GR)_ = 0.0156). There was a strong correlation between pa-*BI-1* expression and pa-*GR* expression following the gene knockdown (lm, GR∼BI-1, p = 0.0036, r^2^ = 0.48), indicating the inhibition of pa-*BI-1* expression leads to a decrease in pa-*GR* expression.

To exclude non-specific effects of the siRNA treatment, we analyzed the correlation of pa-*BI-1* expression with 4 genes (pa-*HSP70*, pa-*Bcl-2*, pa-*BAK*, pa-*BAX*) previously found to be involved in coral bleaching^41, 50, 51^ and which follow a similar expression patterns as pa-*BI-1* in PC and NPC corals upon heat stress, and with pa-*NFKBI* (NFkB inhibitor) that was differentially expressed (^41^, Fig S3A). *pa-BAX* and *pa-Bcl-2* showed strong correlation with *pa-BI-1* gene expression in PC corals (Fig S3B) (r_(Pearson)_ = 0.856 and 0.726; p_(lm)_ < 0.001) and a weaker but still considerable correlation in NPC corals (r_(Pearson)_ = 0.642 and 0.531; p_(lm)_ < 0.001). *pa-BAK* and *pa-HSP70* showed a moderate correlation with pa-BI-1 in PC corals (r_(Pearson)_ = 0.446 and 0.438; p_(lm)_ = 0.013 and 0.0023) but – as expected – the expression of *pa-NFKBI* did not correlate with the expression of *pa-BI-1* in any corals (r_(Pearson)_ = 0.27 and 0.203 (for PC and NPC corals, respectively); p_(lm)_ = 0.12 and 0.18).

We hypothesized that if the siRNA treatment is not specific to the *pa-BI-1* gene or impacts the whole bleaching pathway, we would observe a shift in multiple genes involved in the coral bleaching cascade. Upon siBI-1 treatment, the expression of none of these genes changed significantly (Fig S3B), supporting the specificity of the siRNA treatment to siBI-1 gene expression.

### Decrease in glutathione reductase gene expression results in a decrease in enzyme activity

Changes in gene expression are not always directly mirrored in the protein level and/or activity ^42^, so we measured the activity of glutathione reductase in coral host tissue after siBI-1 gene knockdown. The decrease in pa-*GR* expression after 3 hours of heat stress is followed by a significant decrease in the enzymatic activity at 24 hours post stress (Paired t-test, p_(activity)_ = 0.0391, Fig. 4E). There was a rapid decrease in the glutathione reductase activity in 5 corals, and a very slight increase in 2 corals (Fig. 4E; dotted lines connecting siNTC-siBI-1 pairs), suggesting that decreased pa-*GR* expression does result in decreased pa-GR activity, but there may be genotype-specific effects.

### Corals with inhibited expression of pa-BI-1 are more prone to oxidative DNA damage

To examine the connection between antioxidant system and oxidative DNA damage in heat-stressed corals, we analyzed the level of oxidative DNA damage using an 8-OHdG marker in corals with pa-*BI-1* knockdown and subsequent decrease in glutathione reductase expression and activity. siBI-1 treated corals accumulated significantly more oxidized guanines in DNA during acute heat stress (Fig. 4E; 24 hours at 32 °C, Paired t-test, p = 0.0181) when compared to the PC corals treated with control siRNA. This observation supports the hypothesis that during heat stress, corals use antioxidants to protect important cellular structures from oxidative damage.

### An antioxidant responsive element (ARE) lies within the promotor of coral glutathione reductase gene

In mammals, BI-1 can activate the Nrf2 transcription factor which regulates gene expressions through the *cis*-acting elements called antioxidant responsive elements (ARE) ^45, 52^ in various Nrf2 target genes’ promoters. Nrf2 gene has not been described in reef-building corals but the *Nematostella vectensis* putative Nrf2 protein sequence (GenBank KU746947.1), returns blast hits (tblastn) for several uncharacterized loci in scleractinian corals (e.g., *Orbicella faveolata* LOC110060612, 93 % query cover, 29.89% identity and 1e-43 E-value, or *Pocillopora damicornis*, LOC113687044, 53% query cover, 30.94% identity and 1e-43 E-value) suggesting a protein with a Nrf2-like function may be present.

We searched the promoter region (within NW_020843386.1) of the glutathione reductase gene (XM_027196629.1) in *Pocillopora damicornis* ^53^ and found an ARE-similar sequence 5‘- TGACTTAGC-3’ ^52, 54^ 557 bp upstream of the predicted beginning of the gene ORF. This ARE was first discovered and described in the promoter of glutathione peroxidase in human liver carcinoma cells (HepG2) at positions −76 and −387 with respect to +1 transcription start site ^54^. This striking resemblance suggests that the Nrf2/ARE pathway may be more evolutionarily conserved than previously thought.

## Discussion

Like most organisms, corals have the ability to acclimatize to environmental change, which reduces the severity of bleaching and mortality during thermal stress ^34–38, 41, 55^. However, this natural phenomenon may be genotype-specific ^39^ and could be lost under future climate-change scenarios ^40^. One of the main obstacles to the application of this strategy for conservation ^7, 8, 33^ is our limited understanding of molecular and cellular mechanisms behind acclimatization and/or adaptation to increased temperatures. Here we show that preconditioning-based acclimatization is mediated by the interaction of the pro-life gene *BI-1* and the antioxidant response, which impacts fitness via cellular phenotypes such as DNA damage.

Transcriptomic data from acclimatized corals shed light on the main gene families and cellular pathways that play a role in the bleaching process, but functional studies have been largely missing in reef-building corals ^34, 35, 37, 56^. Recently, we showed that preconditioning in *Pocillopora acuta* leads to improved thermal tolerance due to modulations in the programmed cell death pathway (PCD), most likely via autophagy/symbiophagy ^41^. However, the primary signals or molecular consequences of such a prolonged symbiosis maintenance under thermal stress are unclear. During heat stress, the coral host and symbionts release increased reactive oxygen species (ROS) ^11, 12, 14, 15, 19, 25, 57^ which can activate an array of regulatory pathways, often depending on the level of ROS accumulation ^58, 59^. For example, in model animals, low doses of ROS activate cell survival signaling pathways such as the unfolded protein response (UPR) or Nrf2, while high doses of ROS activate PCD ^59^. We thus hypothesized that after preconditioning, the level of ROS signaling molecules in heat-stressed corals was reduced, which resulted in changes in PCD signaling.

We find that preconditioned (PC) corals with higher tolerance to thermal stress and reduced bleaching rate ^41^ have higher activity of glutathione reductase, but not peroxide-scavenging enzymes, in the host tissue relative to non-preconditioned (NPC) corals. Gene expression analysis suggests that the observed increase in glutathione reductase activity derives from the host but cannot be explained simply and entirely by the expression rate. The levels of mRNA do not always correlate with cellular protein levels and the relationship between the two strongly varies during dynamic transitions such as short-term acclimatization (reviewed in ^42^), a pattern which was previously observed in heat-stressed *Symbiodiniaceae* ^60^ While posttranscriptional and posttranslational modifications may positively impact the activity, turnover rate, or localization of the antioxidant in PC corals in ambient conditions, our results suggest the rapid induction of pa-*GR* gene expression in PC corals is potentially a major contributor to the observed activity differences upon acute heat stress.

Glutathione is a non-enzymatic antioxidant that exists in reduced (GSH) and oxidized (GSSG) form in the cell (reviewed in ^20^). GSH neutralizes ROS while being oxidized to glutathione disulfide (GSSG); this oxidized state is converted back to the reduced state by glutathione reductase (GR). Under normal conditions, over 90 % of the glutathione pool is maintained as GSH by GR activity, so GR is directly responsible for maintaining the reducing environment of the cell and for ROS scavenging. The level of GSH and/or the activity of enzymes involved in the glutathione redox cycle are inversely associated with the oxidative DNA damage ^61, 62^ ROS-related oxidative DNA damage can stem from heat stress in mice, human, plant and fish cells ^63–68^, and increased DNA damage was observed in coral tissue explants exposed to elevated temperatures or to direct sunshine ^21, 69^. Inversely, heat acclimation prevents accumulation of 8-OHdG markers in blood cells of navy boiler tenders exposed to high heat during work ^70^.,

We tested the level of oxidized guanine species, markers of oxidative DNA damage, in PC and NPC corals after 24 hours of heat stress and found that while NPC corals accumulate these markers, PC corals do not. Previous experiments showed that the addition of the exogeneous antioxidant mannitol can reduce DNA damage in heat-stressed coral cell aggregates ^21^, so we used this antioxidant on a new set of NPC corals to solidify the link between antioxidant system and DNA damage in the whole adult coral organism, demonstrating that the antioxidant system is the major contributor to the differences in DNA damage between PC and NPC corals.

DNA damage also triggers diverse cell death pathways, including autophagy ^71^, supporting our previous results showing preconditioning improves coral thermal tolerance via modulations in the autophagy pathway ^41^. BI-1 (BAX-1 inhibitor), a pro-survival PCD gene involved in the regulation of PCD pathways, was shown to reduce ROS accumulation in vertebrates and to activate Nrf2, which is a transcription factor of various antioxidants, including glutathione reductase ^45, 72^. Interestingly, we saw an upregulation of pa-*BI-1* in PC corals upon heat stress, peaking at 3 h ^73^, which parallels host glutathione reductase expression. We compared the expression rates of pa-*BI-1* and pa-*GR* in PC and NPC corals and strikingly, they are highly correlated in PC corals in the early phase of the heat stress response (< 6hours), but not in NPC corals (Fig. S2). We hypothesize that preconditioning may enable pa-BI-1 to effectively regulate the expression of pa-*GR* through epigenetic modifications of the pa-*GR* regulatory elements. DNA methylation patterns vary between corals living in different environments and is dynamic over time in corals exposed to environmental changes, very likely enabling fine tuning of expression in response to various conditions ^74–77^. Future experiments should investigate this link between preconditioning, epigenetic modifications, and expression of particular genes in corals.

To demonstrate the functional correlation between pa-BI-1 and pa-GR, we developed a protocol for siRNA-mediated gene knockdown in living adult corals. Gene knockdown assays have shown limited efficiency in Cnidarians and have only been used in a few studies, representing a major obstacle for functional studies of these non-model organisms. Existing protocols in cnidarians were not technically feasible to adult reef-building corals, relying on microinjection of juveniles or electroporation of anemones as the delivery technique^47–49, 78^. Other studies using liposomal compounds transferred long dsRNA into anemones or coral larva, where the animal’s RNAi machinery was allowed to process it into siRNA^48, 79, 80^. The skeletal structure of adult reef-building corals prevents these approaches from being effective.

In this experiment, we used processed siRNA designed to specifically target the gene of interest and delivered it to the coral tissue using a lipidic vesicle INTERFERin (Polyplus) which proved to be the most efficient and the only repeatable method of those tested. We first targeted the family of green fluorescent proteins, a standard strategy to validate gene-knockdown in non-model organisms owing to its ease of monitoring success in real time ^47–49^. We were able to achieve a significant downregulation of GFP proteins in the majority of test corals and we hypothesize that coral mucus likely interferes with the siRNA transfection procedure in some individuals, meaning that individuals undergoing siRNA-involved experiments must be first tested for the successful knockdown before further functional analyses take place. Additionally, the knock-down efficiency we achieved allowed to study the function of genes that are upregulated during specific cellular and molecular processes (after the overexpression is knocked down close to the basal level) but may not be applicable for complete loss-of-function studies.

Our siRNA methodology allowed us to functionally connect BI-1 gene expression with the coral antioxidant system by inhibiting pa-*BI-1* overexpression in preconditioned corals during acute heat stress. We manipulated the gene in 8 out of 17 corals, showing decreased pa-*GR* expression leading to a decline in pa-*GR* activity and proving that pa-BI-1 regulates gene expression of pa- GR in preconditioned corals. The decreased expression of pa-*GR* paralleled decreased glutathione reductase antioxidant activity in siBI-1 corals, leading to more oxidative damage than corals treated with a control siRNA (siRNA with no known target in *P. acuta*), supporting the hypothesis that antioxidants prevent cellular oxidative damage in corals.

In vertebrate models, BI-1 was also found to regulate expression of genes coding for antioxidants through Nrf-2 transcription factor ^45, 46^. Although Nrf2 has not been described in reef-building corals, a homolog of the Nrf-2 gene was identified and annotated in the anemone *Nematostella vectensis* (GenBank KU746947.1, ^81^), where the Nrf-2 mediated oxidative stress response pathway was activated in its symbiotic but not apo-symbiotic morph during thermal stress ^82^. Nrf2 regulates the expression of antioxidant genes via binding to the so-called ARE (antioxidant response element) *cis*-elements located upstream of the transcription start site ^52^. In the putative promoter of coral glutathione reductase gene, we found an ARE similar to one described in the promoter of the glutathione peroxidase gene in human liver cells ^54^, suggesting the Nrf2/ARE pathway is conserved in Cnidarians, where it controls the expression of antioxidant genes during environmental stress response. We thus propose this pathway could also connect BI-1 and glutathione reductase in *Pocillopora acuta*.

This work describes cellular and molecular principles of coral symbiosis maintenance under heat stress and how it is modulated during acclimatization, which is a critical ecological pathway to the long-term persistence of coral reefs. We propose that pre-exposure to thermal stress improves the ability of corals to maintain cellular redox homeostasis through BI-1-mediated glutathione reductase overexpression, which prevents the accumulation of oxidative stress markers, avoids the activation of programmed cell death and results in prolongated coral-algal symbiosis. Conversely, heat-stressed corals may experience high ROS, subsequent oxidative stress, activation of cell death pathways, and bleaching. Functional studies contribute to our growing knowledge of the response of organisms to environmental stress and ecosystem trajectories under climate change.

## Materials and Methods

### Collections, experimental setup, and preconditioning

Seven colonies of *P. acuta* were collected in spring 2018 at different sites and depths ranging between 1 to 4m across Kāne‘ohe Bay, Hawai‘i to maximize genetic diversity. Experimental treatments followed in Majerova et al. ^41^. Briefly, corals were fragmented and allocated into preconditioning (PC) and control (NPC) treatments and PC corals were exposed to a 29°C for 72h before returning to ambient while NPC corals were maintained at 26°C (Fig 1A). Fragments were then clipped into 5cm nubbins and reallocated into heat stress treatments. After two weeks at 26°C, corals were exposed to a 32°C treatment or control and sampled at 0, 1, 3, 6, 12, and 24 hours. Fragments were stored immediately at −80°C. Bleaching rate was assessed as the shift in symbiont-to-host signal ratio with time-lapse confocal microscopy (Zeiss LSM-710) as described in Majerova et al.^41^

### Antioxidant activity assays

Samples previously stored at −80°C were homogenized in ice cold extraction buffer (100mM Tris-HCl, 20mM EDTA, pH 7.5) in Qiagen TissueLyser (30s^-1^ for 20s) with acid-washed glass beads (Sigma). The symbiont and host cells were separated with low-speed centrifugation (800g, 5min, 4°C) and the supernatant was then sonicated for 3 mins. Cell debris were pelleted by centrifugation (14,000g, 10min, 4°C) and whole cell protein extract concentration was measured using Qubit Protein Assay Kit (Thermo Fisher). Protein extracts were immediately used for EnzyChrom Catalase Assay Kit and EnzyChrom Glutathione Reductase Assay Kit (BioAssay Systems), respectively. The working protein concentration was optimized prior to the experiment to ensure the measured activity was within the range of kit detection limits. We used 1340ng and 300 ng of the whole cell protein extract in Catalase Assay Kit and Glutathione Reductase Assay Kit, respectively, to normalize the enzymatic activity to total protein. Each analysis included one sample per timepoint per colony (n=7).

The Catalase Assay Kit is not specific to catalase and measures activity of all peroxide-scavenging antioxidants, so we used the EnzyChrom Glutathione Peroxidase Assay Kit (BioAssay Systems) to distinguish activity of different peroxide-scavenging antioxidants. All assays (total protein 2 – 10μg) were below the detection limit of the kit.

### Gene expression analysis

Gene expression was analyzed as described in Majerova et al.^41^. Briefly, RNA was extracted via RiboZol RNA Extraction Reagent (VWR Life Science) with DNase I step between phenol-chloroform extraction and ethanol precipitation. 1μg of RNA was reversely transcribed with High-Capacity cDNA reverse transcription kit (ThermoFisher Scientific). Reverse transcription quantitative PCR (RT-qPCR) reactions were run in 12 μl with PowerUp^TM^ SYBR^TM^ Green Master Mix (Applied Biosystems) for 40 cycles. Efficiency of each primer pair was tested with a dilution curve followed by melting curve analysis. Profiles of four genes for each organism (host and symbiont) under different treatments were compared with Best Keeper software and pa-*EF-1* (elongation factor 1a, *P. acuta*), and s*SAM* (S-adenosyl L-methionine synthetase, *Symbiodinium sp.)* showed the highest expression stability upon heat stress and were chosen as the reference genes. All primer sequences are listed in Table S1. Expression of target genes was calculated relative to the non-preconditioned coral at time 0 with ΔΔCt method.

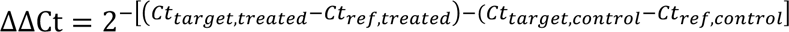

Each analysis included one sample per timepoint per colony (n=7). Sequences of all primers used in this study are listed in Table S1.

### Oxidative DNA damage assay

To assess the impact of oxidative stress on DNA, we analyzed the level of 8-Hydroxy-2’- deoxyguanosine (8-OHdG) in PC and NPC corals as a proxy for cumulative oxidative DNA damage. 2 fragments from 6 corals in each treatment were sampled prior to and after 24 hours of thermal stress (as described above), frozen and kept at −80°C.

We used a custom extraction protocol to extract DNA from host cells after symbiont and host cells were separated as described above. After the low-speed centrifugation, SDS (0.5% final concentration) was added to the supernatant containing host cells and nuclei and incubated at 58°C for 15 min. Then, proteinase K (0.5 mg/ml final concentration) was added and incubated at 58°C for 2h. After incubation, samples were cooled to room temperature and KAc was added to a final concentration 0.5M. Samples were centrifuged (14,000g, 10min, 4°C) and the supernatant was precipitated by isopropanol (1:0.7 ratio). After precipitation, the DNA pellet was resuspended in water and the sample was treated by RNAse A for 15 min. DNA was purified by phenol-chloroform extraction followed by ethanol precipitation (O/N, −20°C) and resuspended in water. 500ng of total genomic DNA in 20μl was denatured at 95°C for 5 min, rapidly cooled down on ice, digested to single nucleotides by 20U of Nuclease P1 (NEB) at 37°C for 30 min and dephosphorylated by 1U of Shrimp Alkaline Phosphatase (NEB) for 30 min at 37°C. All the enzymes were inactivated at 75°C for 10 min. DNA was then diluted 1:5 in 1x Assay buffer and processed according to the kit instructions (DNA Damage Competitive ELISA Kit, Invitrogen). The initial DNA concentration used in the assay was optimized to ensure the resulting values fall within the optimum range of kit detection limits (between Std3 and Std6). We used MyCurveFit.com software to model the standard curve and to predict result values. The analysis was conducted in 2 samples per timepoint per colony using 6 colonies in total. The results from biological replicates were averaged before statistical analysis.

Due to the results indicating that corals with increased glutathione reductase activity are less prone to oxidative DNA damage accumulation, we decided to test the impact of exogenous antioxidants to heat-stressed non-preconditioned corals. For this analysis, a new set of corals was collected from Kāne’ohe Bay in spring 2020 and prepared as describe above. This new set of corals was used for the following mannitol experiment and all the siRNA-involved experiments (see below).

To analyze the impact of exogenous antioxidant to oxidative DNA damage, we treated NPC nubbins with 10mM mannitol ^21^ or a seawater control for the duration of the heat stress experiment (6 NPC and 6 PC corals, 2 nubbins per colony per treatment). All coral fragments were sampled after 24h of acute heat stress (32°C), frozen and kept at −80°C. DNA damage assay was conducted as described above.

### siRNA design

BLOCK-iT™ RNAi Designer (ThermoFisher Scientific) was used to design siRNA for the *P. damicornis* BI-1 mRNA sequence (XM_027189407.1, ^53^) and predicted GFP-like mRNA (XM_027188377.1). In order to silence the most GFP-like genes possible, we aligned all the proposed siRNA sequences to the set of 12 predicted GFP-like proteins we found in the *Pocillopora* transcriptome database in NCBI (XM_027191428.1, XM_027191431.1, XM_027188381.1, XM_027195143.1, XM_027188383.1, XM_027188379.1, XM_027188378.1, XM_027188377.1, XM_027188376.1, XM_027195136.1, XM_027188380.1, XM_027188372.1) and chose the sequence with the highest alignment to the most proteins. For BI-1 silencing, we chose the highest-ranking siRNA sequence containing BCD Tuschl’s patterns and targeting region 194- 212 of the 1437pb long mRNA sequence. Control siRNA (siNTC) was designed to contain the same nucleotide composition as siBI-1 but having no known target in the *P. damicornis* mRNA sequence database. All siRNA molecules were synthesized by Sigma-Aldrich. Sequences of all siRNA molecules are listed in Table S1.

### siRNA-mediated knockdown

PC coral fragments (∼ 3 cm long) were placed into 20 ml cultivation vials, completely submerged into flow-through tanks with ambient temperature seawater and left to acclimatize overnight. The next day, vials were moved into precise temperature-controlled water bath, the seawater was carefully removed without disturbing corals and the siRNA transfection was carried out according to the manufacturer’s instructions (INTERFERin® transfection reagent, Polyplus). siRNA transfection mix (20 µl of 10nM siRNA and 16 µl INTERFERin® reagent in 250 μl 0.2nm filtered seawater) was pipetted directly on the exposed coral and after ∼ 2 minutes, 5 ml of 0.2 nm filtered seawater was added to fully cover the nubbin and coral was incubated at ambient temperature for 6 hours. After this time, corals were fully submerged into a flow-through tank with ambient temperature seawater for two days. For siBI-1 experiment, 48 hours after the beginning of the siRNA transfection, corals were exposed to an acute heat stress (32°C, ramping speed of ∼1°C per 10 mins). At 3 hours post-stress, each coral was sampled for the gene expression analysis and at 24h hours post-stress for glutathione reductase activity and DNA damage assay. This sampling consumed the fragment.

### Confocal microscopy

Corals for the siGFP experiment were microscoped as described in Majerova et al.^41^, with the following adjustments. Each coral nubbin was scanned three times with EC Plan-Neofluar 2.5x/0.075 objective using Z-Stack mode with fixed 12 layers spanning tissue thickness of approximately 0.1 mm. Images were then processed in Fiji software (ImageJ)^83^ using Zprojection set to Sum Slices to display the sum of the symbiont and GFP-derived signals through all the tissue layers. We then measured the signal intensity in 10 circular regions of interest (ROIs) spreading to the whole fragment with exception to polyp mouths, tentacles, and clearly injured tissue (after cutting the nubbin, e.g.) and compared the ratio of GFP signal to the symbiont-derived signal (chlorophyl a) between siGFP and siNTC treated corals.

### Statistical analyses

All statistical analyses were conducted in R. Datasets were tested for normality (histogram plot) and heteroscedascity (Levene test) and normalized as needed using a logarithmic transformation or the *BestNormalize* function in R. We analyzed antioxidant activity after normalizing data to the initial control activity (NPC, time 0) for each colony. We used a generalized mixed model (lmer function, package lme4) with treatment and time as main effects and colony as a random effect with a Tukey post-hoc testing for each timepoint.

We analyzed gene expression after normalizing data to the initial control expression for each colony. We used a generalized mixed model to examine normalized expression of each gene separately, using conditioning treatment and time as main effects and colony as a random effect. We estimated least-squares means and compared between treatments for each timepoint using bonferroni adjusted p-values.

To analyze DNA damage, we ran a two-way ANOVA to compare the level of 8-OHdG (pg per μl DNA) of PC and NPC corals with time and conditioning treatment as main effects and a Tukey HSD for post-hoc analysis. The effect of mannitol treatment to DNA damage was tested by one-way ANOVA with treatment (control, heat stress, heat stress with mannitol) as variable followed by a Tukey post-hoc testing.

We calculated Pearson correlation coefficients and variance explained using linear regression for the relationships between a) gene expression and enzymatic activity and b) BI-1 and glutathione reductase expression.

The effect of siRNA-mediated knockdown was analyzed using paired samples one-tailed T-test for the siGFP experiment (gene expression and microscopy) and paired samples Wilcoxon test on data normalized to control (ambient temperature, non-treated) coral samples for the siBI- 1 experiment. Antioxidant activity data in siBI-1 experiment were also first normalized to control coral samples and then subjected to paired T-test. DNA damage analysis in siRNA-treated corals was analyzed with paired t-test on raw data.

## Data availability statement

All data needed to evaluate the conclusions are present in the manuscript and Supplementary Materials. Data available on request from corresponding author.

## Acknowledgements

We would like to thank Shayle Matsuda, Fiona C. Carey, Filip Blaštík, and Vojtěch Prokůpek for their help with coral collection and siRNA knock-down optimization tests, and Tomáš Buryška for advice on execution of enzymatic assays. We thank the ToBo Lab (HIMB) for access to instruments. We dedicate this manuscript to Ruth Gates, who inspired us to use molecular tools to understand the coral bleaching crisis. This work was funded by the Paul G. Allen Family Foundation, the Annenberg Foundation and the National Science Foundation (IOS-2041401). Corals were collected under SAP-2020-25 to HIMB. This is HIMB contribution xx and SOEST contribution xx.

## Author Contributions

**EM** conceived the experiments, conducted research, analyzed data, and wrote the manuscript.

**CD** analyzed data and wrote the manuscript. Both authors approved the final version.

## Supplementary material

**Figure S1.**
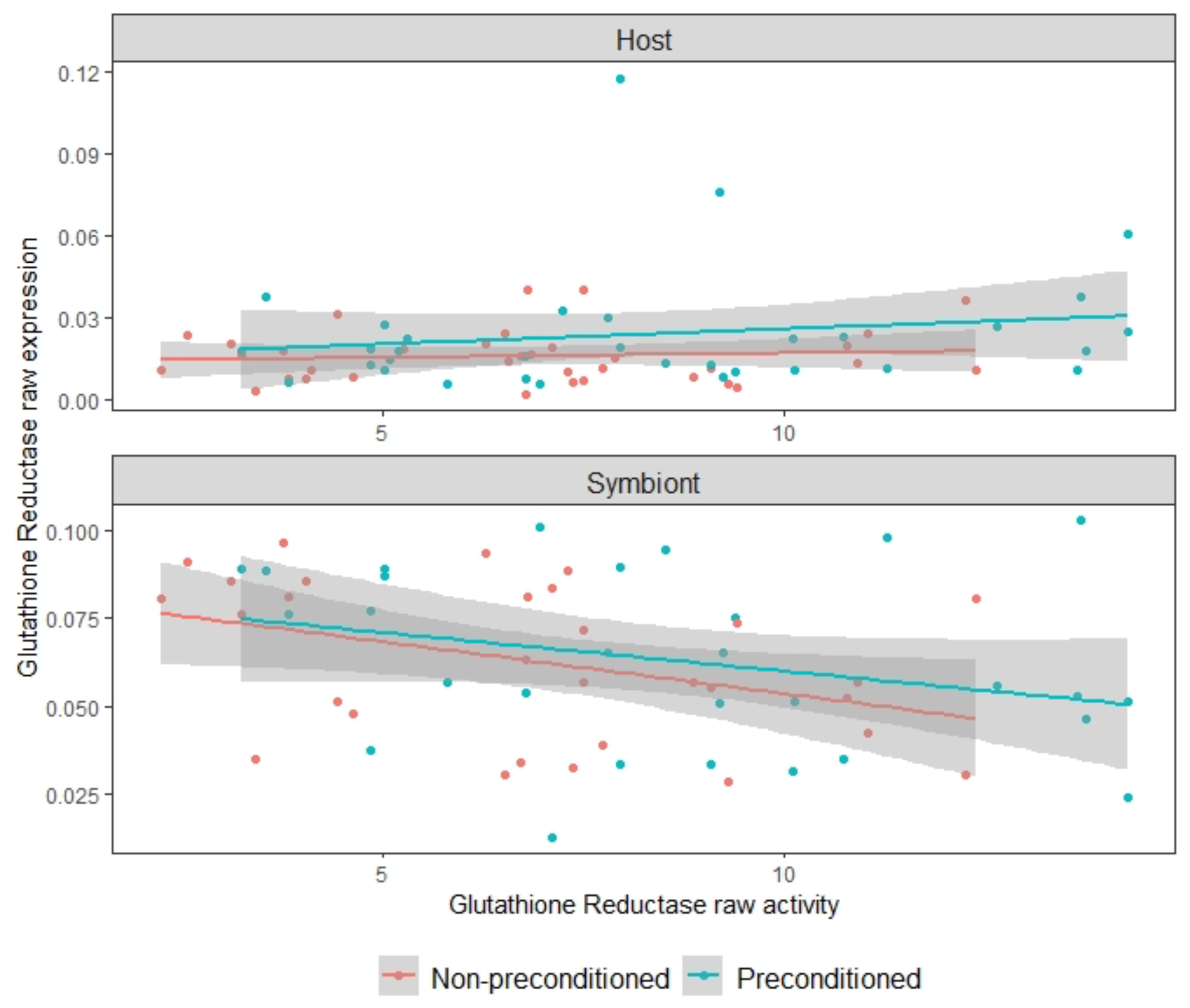
Correlation of glutathione reductase activity and gene expression across all timepoints. There is no apparent correlation between the gene expression in host or symbiont cells with the activity of the enzyme. Raw expression is calculated as 2^-ΔCt^ ;ΔCt = Ct_(target)_ – Ct_(reference),_ raw activity is measured in U/L. n = 6.

**Figure S2.**
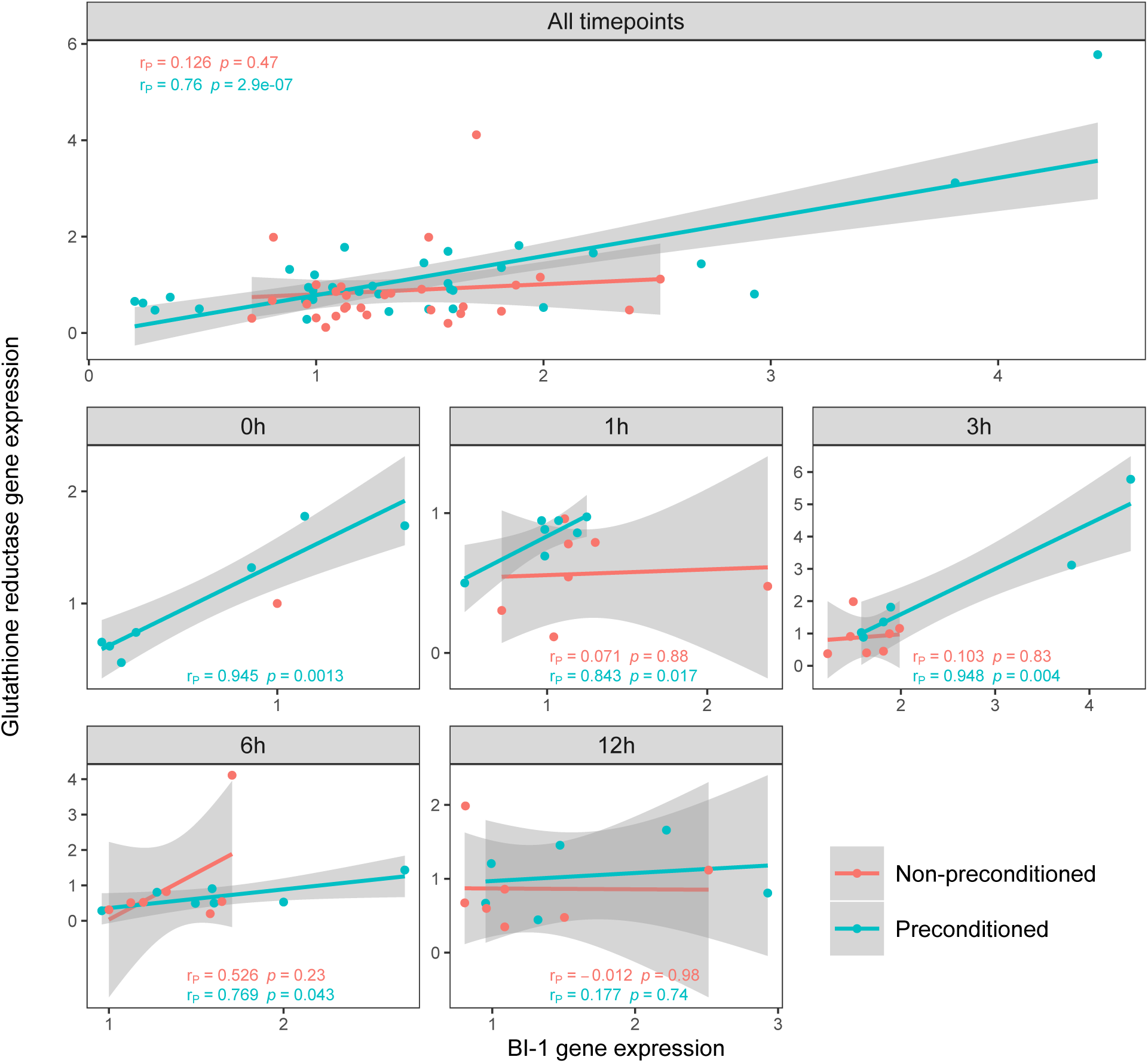
Expressions of BI-1 and glutathione reductase correlate in corals after preconditioning at ambient temperature and during early hours of an acute heat stress. Non-preconditioned (NPC) corals are depicted in red, preconditioned (PC) corals in cyan. All expression levels are normalized to the control coral (NPC at ambient temperature) using 2^-ΔΔCt^ formula. r_P_ stands for Pearson correlation coefficient, p for p-value

**Figure S2.**
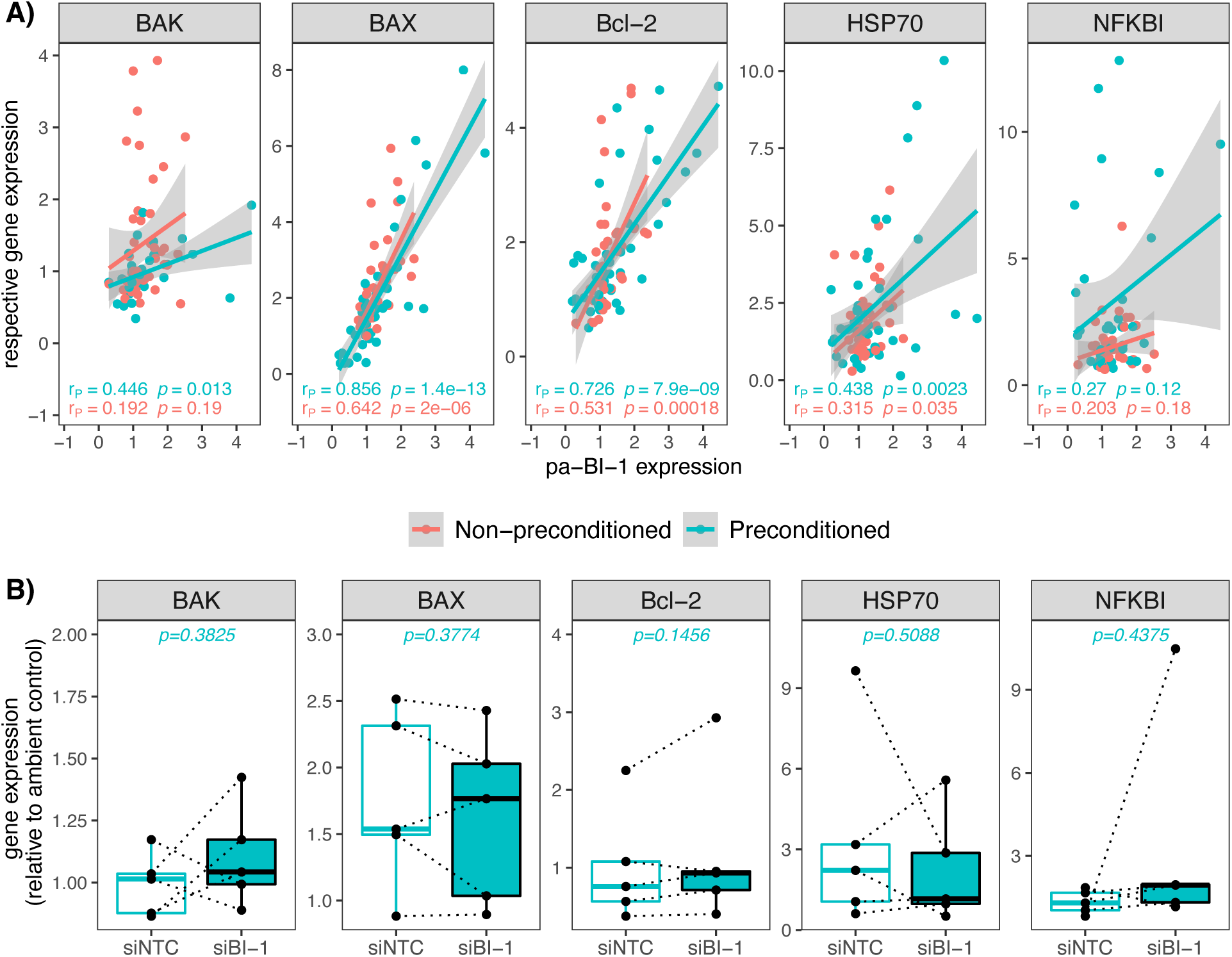
**A) Correlation of of pa-BI-1 gene expression with other genes involved in bleaching pathway.** B) Expression of these genes after pa-BI-1 knockdown. Non-preconditioned (NPC) corals are depicted in red, preconditioned (PC) corals in cyan. All expression levels are normalized to the control coral (NPC at ambient temperature) using 2^-ΔΔCt^ formula. r_P_ stands for Pearson correlation coefficient, p for p-value. The boxplots show median of the data, first and third quartile and respective datapoints.

**Table S1.**
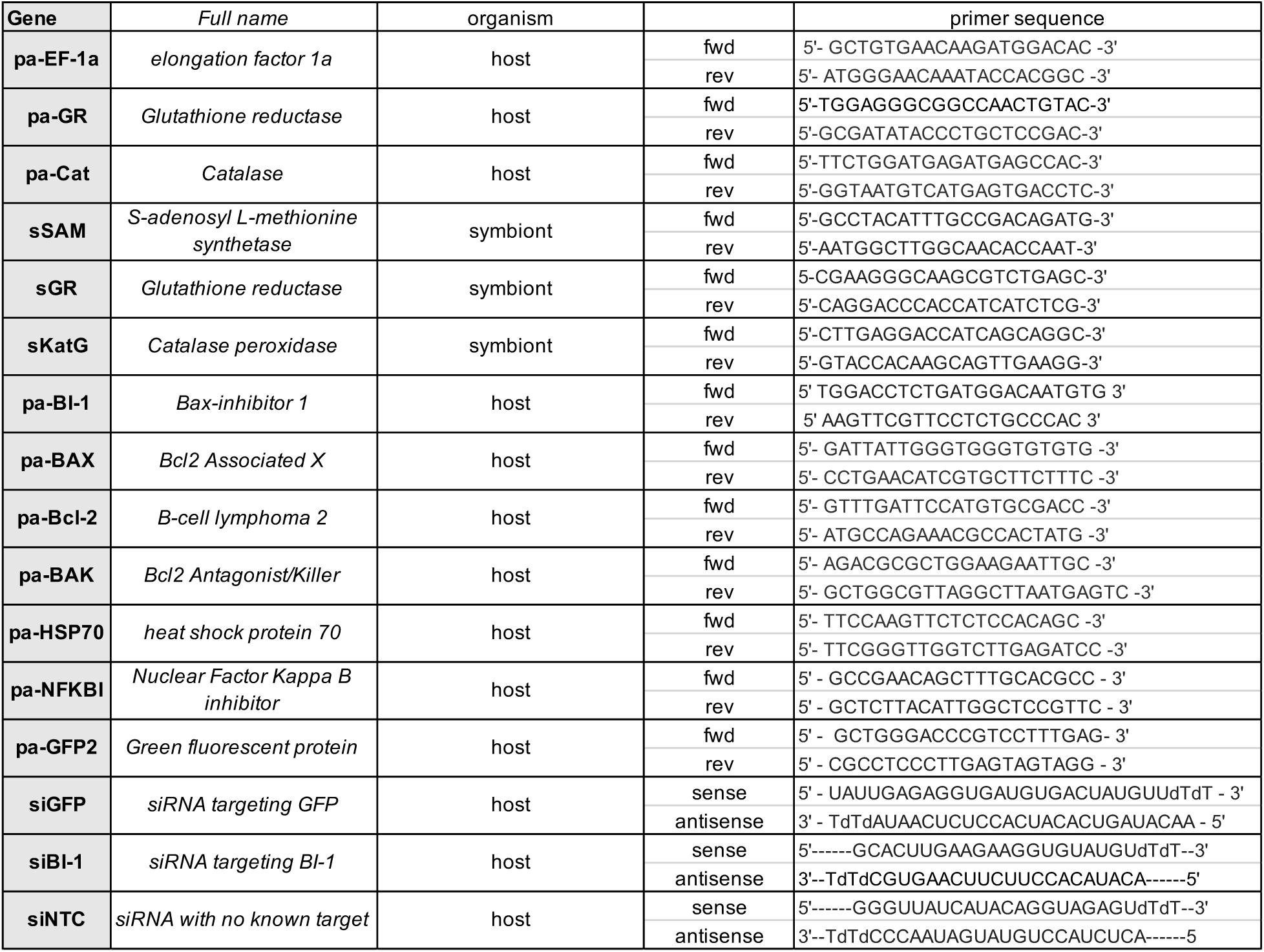
List of primers and siRNAs used in the study.

