## Supplementary material for "Thermal preconditioning in a reef-building coral alleviates oxidative damage through a BI-1 mediated antioxidant response"

Figure S1

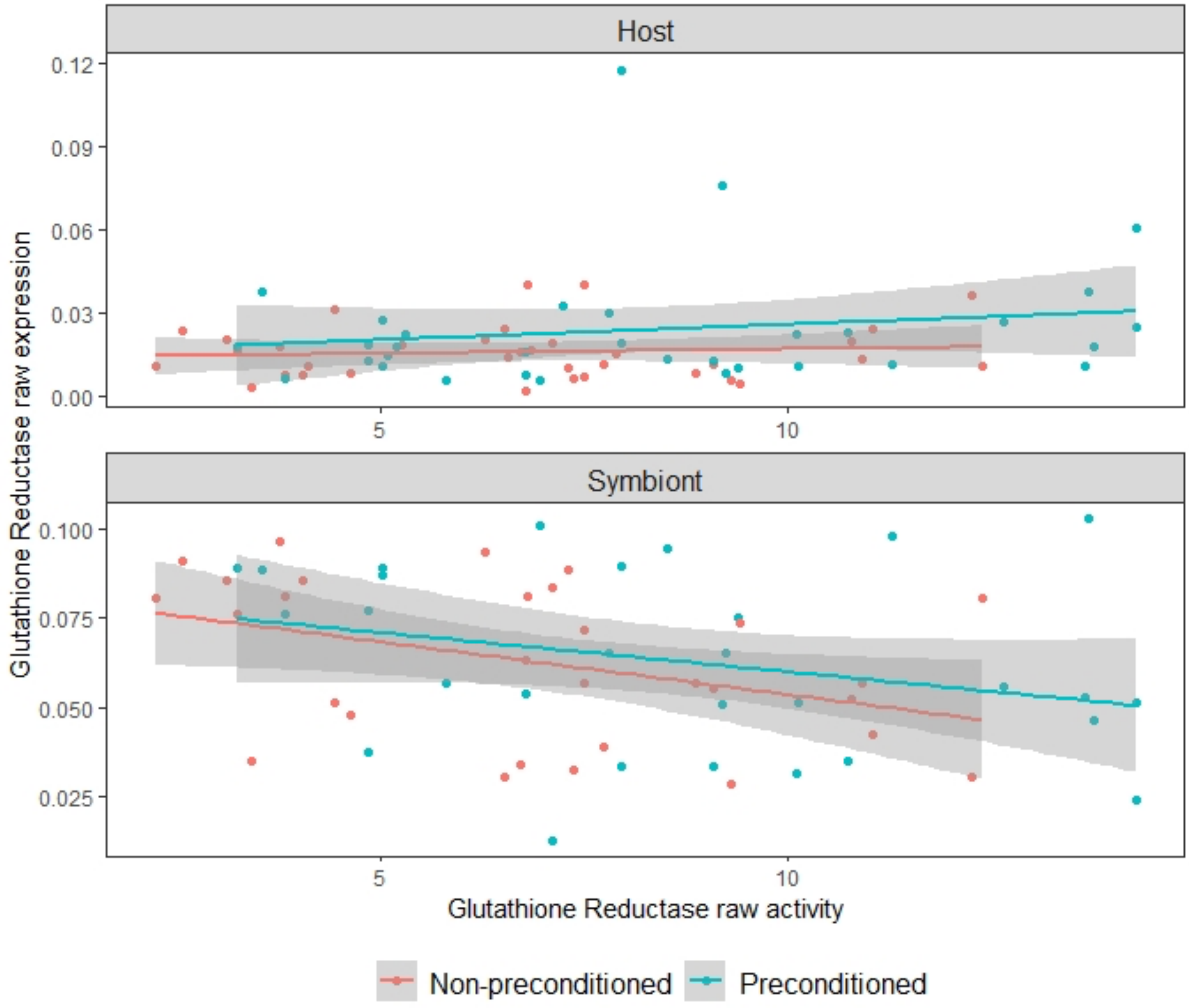


**Figure S1 – Correlation of glutathione reductase activity and gene expression across all timepoints.** There is no apparent correlation between the gene expression in host or symbiont cells with the activity of the enzyme. Raw expression is calculated as $2^{-\Delta Ct}$ ;$\Delta Ct = Ct_{(target)} - Ct_{(reference)}$, raw activity is measured in U/L. n = 6.

Figure S2

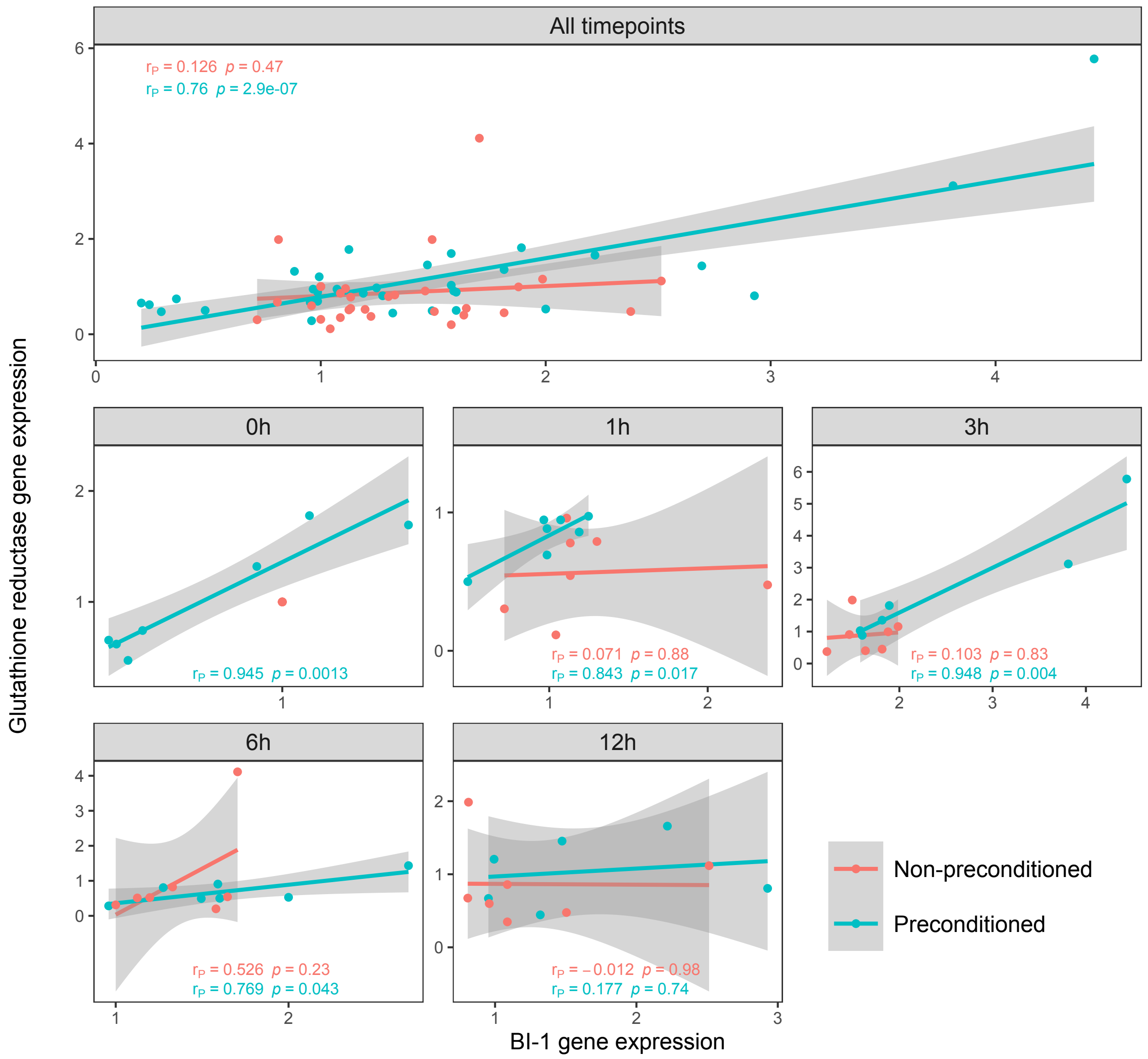


**Figure S2 Expressions of BI-1 and glutathione reductase correlate in corals after preconditioning at ambient temperature and during early hours of an acute heat stress**. Non-preconditioned (NPC) corals are depicted in red, preconditioned (PC) corals in cyan. All expression levels are normalized to the control coral (NPC at ambient temperature) using $2^{-\Delta\Delta Ct}$ formula. $r_P$ stands for Pearson correlation coefficient, p for p-value

Figure S3

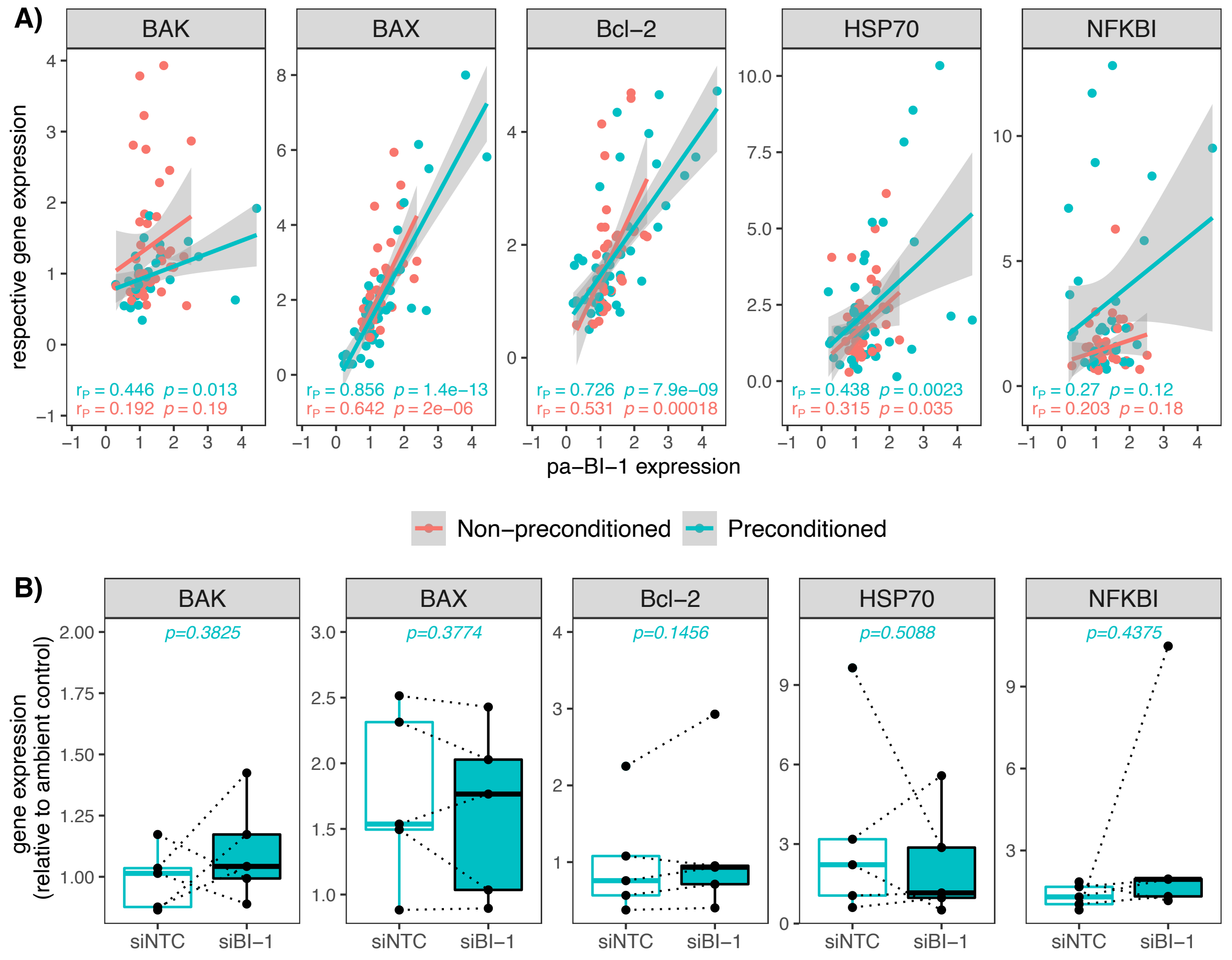


**Figure S2 A) Correlation of of pa-BI-1 gene expression with other genes involved in bleaching pathway.** B) Expression of these genes after pa-BI-1 knockdown. Non-preconditioned (NPC) corals are depicted in red, preconditioned (PC) corals in cyan. All expression levels are normalized to the control coral (NPC at ambient temperature) using $2^{-\Delta\Delta Ct}$ formula. $r_P$ stands for Pearson correlation coefficient, p for p-value. The boxplots show median of the data, first and third quartile and respective datapoints.

Table S1

| Gene | Full name | organism | | primer sequence |
|---|---|---|---|---|
| **pa-EF-1a** | *elongation factor 1a* | host | fwd | 5'- GCTGTGAACAAGATGGACAC -3' |
| | | | rev | 5'- ATGGGAACAAATACCACGGC -3' |
| **pa-GR** | *Glutathione reductase* | host | fwd | 5'-TGGAGGGCGGCCAACTGTAC-3' |
| | | | rev | 5'-GCGATATACCCTGCTCCGAC-3' |
| **pa-Cat** | *Catalase* | host | fwd | 5'-TTCTGGATGAGATGAGCCAC-3' |
| | | | rev | 5'-GGTAATGTCATGAGTGACCTC-3' |
| **sSAM** | *S-adenosyl L-methionine synthetase* | symbiont | fwd | 5'-GCCTACATTTGCCGACAGATG-3' |
| | | | rev | 5'-AATGGCTTGGCAACACCAAT-3' |
| **sGR** | *Glutathione reductase* | symbiont | fwd | 5-CGAAGGGCAAGCGTCTGAGC-3' |
| | | | rev | 5'-CAGGACCCACCATCATCTCG-3' |
| **sKatG** | *Catalase peroxidase* | symbiont | fwd | 5'-CTTGAGGACCATCAGCAGGC-3' |
| | | | rev | 5'-GTACCACAAGCAGTTGAAGG-3' |
| **pa-BI-1** | *Bax-inhibitor 1* | host | fwd | 5' TGGACCTCTGATGGACAATGTG 3' |
| | | | rev | 5' AAGTTCGTTCCTCTGCCCAC 3' |
| **pa-BAX** | *Bcl2 Associated X* | host | fwd | 5'- GATTATTGGGTGGGTGTGTG -3' |
| | | | rev | 5'- CCTGAACATCGTGCTTCTTTC -3' |
| **pa-Bcl-2** | *B-cell lymphoma 2* | host | fwd | 5'- GTTTGATTCCATGTGCGACC -3' |
| | | | rev | 5'- ATGCCAGAAACGCCACTATG -3' |
| **pa-BAK** | *Bcl2 Antagonist/Killer* | host | fwd | 5'- AGACGCGCTGGAAGAATTGC -3' |
| | | | rev | 5'- GCTGGCGTTAGGCTTAATGAGTC -3' |
| **pa-HSP70** | *heat shock protein 70* | host | fwd | 5'- TTCCAAGTTCTCTCCACAGC -3' |
| | | | rev | 5'- TTCGGGTTGGTCTTGAGATCC -3' |
| **pa-NFKBI** | *Nuclear Factor Kappa B inhibitor* | host | fwd | 5' - GCCGAACAGCTTTGCACGCC - 3' |
| | | | rev | 5' - GCTCTTACATTGGCTCCGTTC - 3' |
| **pa-GFP2** | *Green fluorescent protein* | host | fwd | 5' - GCTGGGACCCGTCCTTTGAG- 3' |
| | | | rev | 5' - CGCCTCCCTTGAGTAGTAGG - 3' |
| **siGFP** | *siRNA targeting GFP* | host | sense | 5' - UAUUGAGAGGUGAUGUGACUAUGUUdTdT - 3' |
| | | | antisense | 3' - TdTdAUAACUCUCCACUACACUGAUACAA - 5' |
| **siBI-1** | *siRNA targeting BI-1* | host | sense | 5'------GCACUUGAAGAAGGUGUAUGUdTdT--3' |
|  |  |  | antisense | 3'--TdTdCGUGAACUUCUUCCACAUACA------5' |
| **siNTC** | *siRNA with no known target* | host | sense | 5'------GGGUUAUCAUACAGGUAGAGUdTdT--3' |
|  |  |  | antisense | 3'--TdTdCCCAAUAGUAUGUCCAUCUCA------5 |

**Table S1 – List of primers and siRNAs used in the study**
